# Complete spatiotemporal quantification of cardiac motion in mice through enhanced acquisition and super-resolution reconstruction

**DOI:** 10.1101/2024.05.31.596322

**Authors:** Tanmay Mukherjee, Maziyar Keshavarzian, Elizabeth M. Fugate, Vahid Naeini, Amr Darwish, Jacques Ohayon, Kyle J. Myers, Dipan J. Shah, Diana Lindquist, Sakthivel Sadayappan, Roderic I. Pettigrew, Reza Avazmohammadi

## Abstract

The quantification of cardiac motion using cardiac magnetic resonance imaging (CMR) has shown promise as an early-stage marker for cardiovascular diseases. Despite the growing popularity of CMR-based myocardial strain calculations, measures of complete spatiotemporal strains (i.e., three-dimensional strains over the cardiac cycle) remain elusive. Complete spatiotemporal strain calculations are primarily hampered by poor spatial resolution, with the rapid motion of the cardiac wall also challenging the reproducibility of such strains. We hypothesize that a super-resolution reconstruction (SRR) framework that leverages combined image acquisitions at multiple orientations will enhance the reproducibility of complete spatiotemporal strain estimation. Two sets of CMR acquisitions were obtained for five wild-type mice, combining short-axis scans with radial and orthogonal long-axis scans. Super-resolution reconstruction, integrated with tissue classification, was performed to generate full four-dimensional (4D) images. The resulting enhanced and full 4D images enabled complete quantification of the motion in terms of 4D myocardial strains. Additionally, the effects of SRR in improving accurate strain measurements were evaluated using an in-silico heart phantom. The SRR framework revealed near isotropic spatial resolution, high structural similarity, and minimal loss of contrast, which led to overall improvements in strain accuracy. In essence, a comprehensive methodology was generated to quantify complete and reproducible myocardial deformation, aiding in the much-needed standardization of complete spatiotemporal strain calculations.

## 1 Introduction

Functional assessments of the left ventricle (LV) have become crucial in the diagnosis of structural heart diseases [1]. Organ-level measurements, such as LV ejection fraction (EF) and end-diastolic volume, are key determinants of systolic and diastolic dysfunction [1, 2, 3]. However, the diagnostic and prognostic efficacy of these “global” indices is challenged by the multi-faceted nature of structural heart diseases. These metrics lag as indicators for most cases of early-stage pathology, with additional investigations often required for risk stratification [4, 5]. The analysis and subsequent characterization of myocardial strains are one such investigation used to complement global metrics. The image-based evaluation of myocardial deformation through global longitudinal strains (GLS) has been shown to be a stronger determinant of cardiac diseases at an early stage in comparison with traditional volumetric metrics [6, 7]. Anomalies in GLS have been linked to cellular dysfunction [8, 9]. Specifically, reduced GLS is suggestive of myofiber-level dysfunction despite the absence of abnormalities in conventional biomarkers in the early stages [10, 11, 12]. Despite their merits, such organ-level strain measurements fail to quantify the regional characteristics of LV function accurately. The LV exhibits complex four-dimensional (4D) kinematics, i.e., full three-dimensional (3D) deformation over the cardiac cycle with remarkable transmural variations, thus confounding the accuracy of regional strain calculations and effectuating variability [7, 13]. The characterization of continuous 4D regional deformation offers excellent potential in establishing a complete LV structure-function relationship. Despite advances in medical imaging, the assessment of 4D continuous myocardial deformation remains vastly under-explored.

One of the most commonly used strain estimations to date is two-dimensional (2D) speckle tracking echocardiography (STE) [14]. While automatic segmentation methods have improved the delineation of myocardial features for three-dimensional (3D) STE, variability in strain estimation is significant due to poor tissue contrast and low spatial and temporal resolution [15]. The exceptional soft tissue contrast offered by cardiac magnetic resonance imaging (CMR) has presented itself as an improved alternative to STE. Tissue tagging methods using specialized image sequences have become the most validated method for strain estimation [16, 17]. However, the complex protocols involved in tag deposition and tracking may pose challenges, especially when 4D strain calculations are of interest [18]. Alternatively, studies on strain calculation using feature-tracking (FT-CMR) methodologies have been implemented using standard CMR scans where cardiac motion is computed via interrogation windows in stacks of short-axis (SA) images. Interpolation schemes can be used to derive spatially 3D strains by extrapolating strains estimated at the epi- and endocardium [19]. However, a single long-axis (LA) slice may not be sufficient to estimate regional 3D strains accurately due to the presence of significant combined torsional and longitudinal motion in the heart. Indeed, although we have previously shown that stacks of short-axis (SA) scans can be potentially used to estimate 4D myocardial strains using image registration [20], a spatially dense stack of SA scans is expected to be needed to describe LA deformation accurately. In addition, acquiring high-resolution (HR) images by prolonging the overall image acquisition time is considered to be infeasible due to patient discomfort. Overall, we suggest that strain accuracy and reproducibility can be improved through super-resolution reconstruction (SRR) using a feasible multi-plane image acquisition strategy.

The SRR of HR images using a series or stream of low-resolution (LR) frames has been previously studied [21]. The ability to choose scanning planes at various orientations enables the application of SRR methods to CMR. A host of scans can be acquired along different orientations either through rotation or translation about an axis. Multiple frames in a video or angular projections of images can be used to increase the spatial distribution of pixels without deteriorating the temporal resolution [22, 23]. The first application of SRR to improve MR resolution was reported in 1997, [24, 25] and to date, SRR has been used to enhance spatial resolution in various modalities ranging from functional MRI [26] to positron emission tomography [27, 28]. While, for the most part, the focus has been on relatively static organs such as the brain [27, 29, 30], the application of SRR in CMR is growing in prevalence [31, 32]. Odille et al. [33] suggested a slice-to-volume reconstruction approach which involved an ad-hoc modification of LR images during acquisition, and [31] used combined acquisitions of SA and LA views of the LV in their reconstruction methodology. A consistent challenge in CMR images is the rapid motion and consequent finite deformation of the myocardial wall during the cardiac cycle. Example-based [34] and deep-learning-based [35] super-resolution methods have been suggested in combating frame registration errors due to high-speed cardiac motion. Investigations have revealed significant improvements in image quality which ultimately reduced the total CMR image acquisition times. An impediment in traditional machine learning- (ML-) based SRR frameworks is the requirement of an extensive repository of training data consisting of HR imaging. The improvement of image reconstruction and subsequent reduction in anisotropy [34, 35] offers significant potential in the complete quantification of 4D myocardial motion. More recent studies have implemented semi- and unsupervised learning methods to create SR images without the need for HR images in training the ML model [36, 37]. Despite the promise of SRR in improving the evaluation of cardiac motion, its application remains primarily limited to improving image quality [31, 35]. However, Xia et al. [36] remarked upon the potential benefits of SRR in myocardial strain analysis and suggested that a combination of poor spatiotemporal resolution and signal-to-noise ratio (SNR) must be addressed to reduce variability in strain calculations. Therefore, there is a need for advancing SRR techniques to improve the reproducibility of the complete spatiotemporal myocardial strains, herein referred to as 4D strains.

In this work, we propose an SRR framework to improve the characterization of 4D myocardial strains using standard cine CMR imaging. The SRR framework combines sets of arbitrarily aligned anisotropic LR scans to produce user-defined 3D HR pixel sets or super-resolution (SR) images using scattered data interpolation. Our method utilizes the classical SRR problem, and by classifying pixels in each LR scan based on tissue type, we improve the spatial resolution in distinct regions of the LV. We hypothesize that the reproducibility of complete spatiotemporal strain calculations can be promoted through the improvement of the distinction between each region, i.e., tissue class. The framework is divided into three modules: (i) the classification module, (ii) the interpolation module, and (iii) the registration module (Fig. 1). Following the tissue classification and interpolation, a non-rigid image registration algorithm was used to calculate pixel displacements and derive the 3D Green-Lagrange strain tensor for the duration of a single cardiac cycle. We evaluated the performance of our proposed framework using CMR images of mice because of their high cardiac metabolism rates and fast myocardial motion, therefore facilitating a rigorous experimental setting to test the efficiency of the SRR framework. Additionally, we utilized a novel *in silico* phantom to compare the accuracy of strain calculations using both conventional and super-resolution approaches. These calculations using SR-generated models extend upon current standards of strain estimation, and a complete 4D map with regional and transmural variations was quantified.

**Fig. 1.**
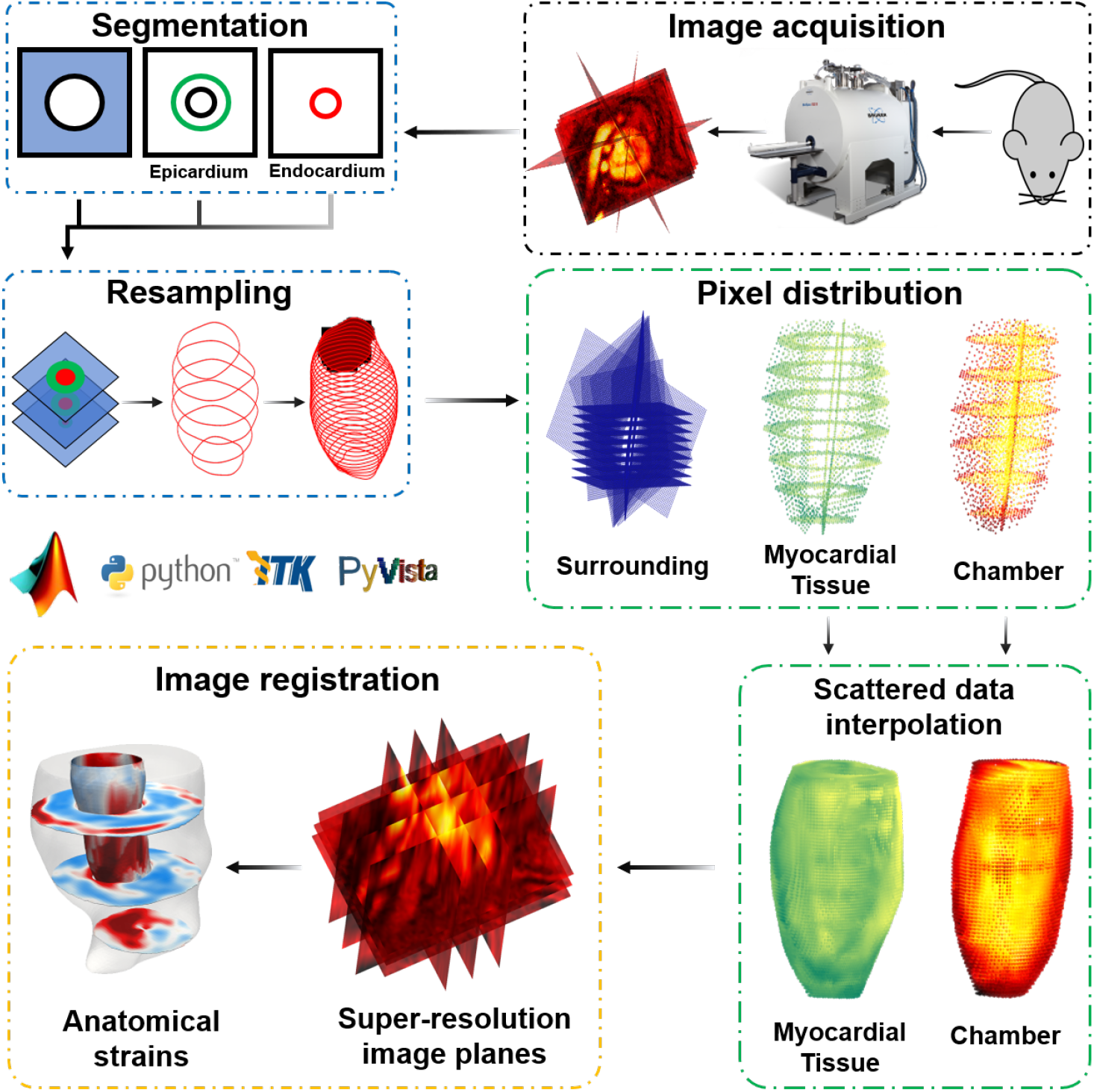
The architecture of the super-resolution reconstruction (SRR) in the cardiac magnetic resonance (CMR) framework consists of (1) image acquisition, (2) SRR, (3) non-rigid image registration, and (4) anatomical strain calculations. 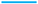 classification module, 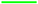 interpolation module, and 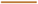 registration module.

## 2 Methods

### 2.1 Development of SRR in CMR

#### 2.1.1 Cardiac tissue classification

The reconstruction of an HR image from sets of LR images can be formulated as an inverse problem. The LR images can be taken as a set of observations( {**y**= **y**_1_,**y**_2_, …**y**_n_}), and the HR image (**x**), is the ideal undegraded discrete representation of the real scene. Each representation can be formulated by combining **x** and random noise, denoted by **n**. Specifically, linear additive Gaussian noise can be assumed when the SNR is greater than 3 for CMR images [38], to form the vectorial representation as:

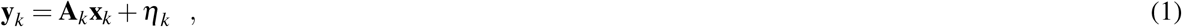

where **A**_k_ describes the image acquisition process for the k^th^ observation. Each observation is a subsampled version of **x**, which is subject to various imaging conditions – namely geometric warping, **W** and blurring, **B**. We can decompose the acquisition model and define **A**_k_ = **D**_k_**B**_k_**W**_k_ through assuming linear space-invariant blurring and linearity in the geometric transform, and downsampling **D** [30, 31]. Thus, for a given set of observations (**y**), an optimization problem is posed to calculate the undegraded HR image (**x**) as:

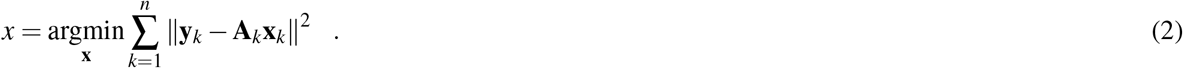

This classical SRR problem can be divided into three steps: (i) registration of the LR images, (ii) interpolation onto a structured or unstructured HR grid, and (iii) deconvolution to remove noise and blur [22]. However, the implementation of this model for CMR images is challenged by the cyclical movement of the cardiac muscle and sharp differences in contrast between the blood pool and the myocardial tissue. To address this limitation, each LR observation was divided into three distinct regions, and an optimization problem was posed to yield the undegraded HR observation. Here, as the focus of the study was cardiac tissue, the regions were defined based on mean gray level intensity into (1) the blood pool, (2) myocardial tissue, and (3) the surroundings. Our objective was to categorize each tissue class (*j*), assign pixels from the image set of the LR acquisitions, and generate query points over a global Euclidean space in the HR grid as:

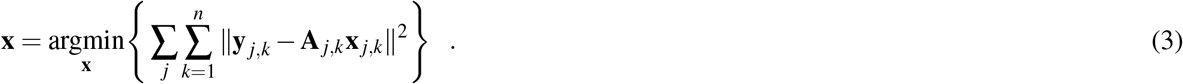

These regions corresponding to the different tissue classes were used to isolate region-specific pixels via a combination of (a) contours drawn on 2D images and (b) information describing surfaces in a structured 3D space. Contours were drawn semi-automatically [39] segment the endocardium and epicardium using Segment version 3.0 R8531 [40]. Subsequently. pixels were assigned to various regions of interest by converting the point clouds into convex hulls and combining the 2D contours into 3D surfaces. These 2D contours were resampled longitudinally through Delaunay triangulation and smoothed through the application of radial basis functions to mitigate the effects of pixel spacing anisotropy on the segmentation quality in 3D reconstruction.

#### 2.1.2 Building tissue class-specific interpolants

The three distinct HR regions were combined by using a priori information about the mean gray level intensity of the region. Subsequently, a regularization term to the right-hand side of the modified SR equation (Equ. 2) allowed the implementation of a priori knowledge about the solution and promoted the smoothness of image gradients at the interface of each region as:

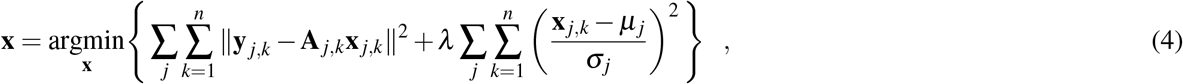

where *µ*_*j*_ and *σ*_*j*_ represent the mean and standard deviation gray level intensity of each *j*, respectively, and *λ* represents the regularization parameter. Thus, deviations from the mean gray level intensity are penalized, which is imperative in improving image registration. The optimization problem (Equ. 4) was used to form region-specific or tissue-class-specific interpolants to solve for **A**_*k*_. The segmented LR images, along with the information describing the desired HR scan, were passed on to the interpolation module. Since images are projected onto either the XY (SA) or the XZ (LA) plane, the coordinates describing the pixel intensities of each image were obtained by accessing the position and direction cosines from the DICOM metadata. The target vector describing k^th^ observation of an LR image was thus obtained, and the rotation matrix was derived using Rodrigues’ rotation formula as:

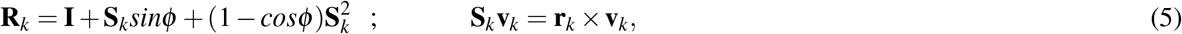

where **R** is the rotation matrix that uses the skew-symmetric transformation matrix *S* obtained from the cross product of the vector **v** and the axis of rotation **r** to rotate the image by an angle *φ*, and **I** is the identity matrix. The rotated image was translated using the position metadata, and the coordinate space describing all observations of the LR images was obtained. Subsequently, the HR scan was allowed to be defined as a structured grid via a global interpolation scheme. Given the anisotropy in pixel distribution and to accommodate arbitrary orientations of LR images, scattered data interpolation was used to estimate **A**_*k*_. The global interpolant consisted of individual regional coordinates interpolated based on their tissue classes. The module allows two regional and global interpolation schemes, namely, (a) *linear*, wherein linear interpolation is used for reconstruction, and (b) *natural* or *pchip*, wherein the HR grid for each tissue class is defined using Delaunay triangulation-inspired piecewise cubic Hermitian polynomials [41]. Each projection of the LR image was evaluated systematically to reduce blurring, and gaps in space were filled over the pre-defined HR grid using the global interpolation scheme, thus yielding a stack of SR images.

#### 2.1.3 Calculating time-course cartesian displacements

The region-specific pixels and a combined package comprising parameters such as the resampling ratios, the reference image index, and the segmented contours were passed on to the registration module. A diffeomorphic demons algorithm [42, 43] was implemented to perform non-rigid image registration. The algorithm aligns a moving image ℳ to the fixed image ℱ through the minimization of the global energy function, **W** as:

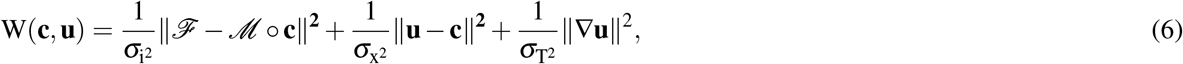

where *σ*_i_, *σ*_x_, and *σ*_T_ are the noise intensity, spatial uncertainty, and regularization factor, respectively, and **u** and **c** denote the parametric and non-parametric spatial transformations, respectively. The registration was performed over three pyramid levels with 500, 400, and 300 iterations.

#### 2.1.4 Estimating anatomical strains

The image registration-derived pixel displacements in the Cartesian frame of reference were used to calculate the Green-Lagrange strain tensor, **E**. Strains were estimated between each reference frame (**x**) and the target frame (**X**) through the deformation gradient tensor, **F**, as:

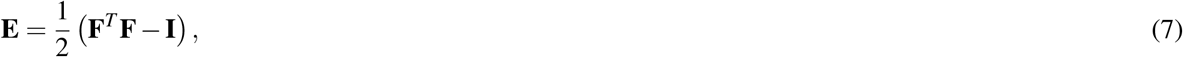

The gradient **F** was calculated as the propagation of the deformation gradient between two consecutive load increments (**F**_i_) such that,

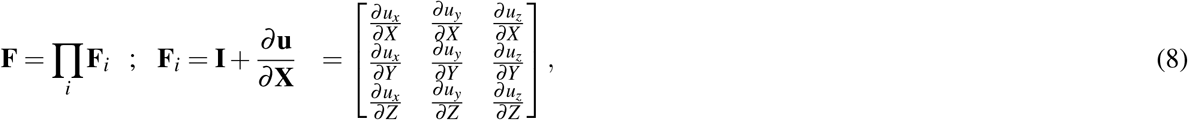

where u_x_, u_y_, and u_z_ are the displacements in the x,y, and z directions between two consecutive images. The resulting Cartesian strains were converted to the widely used anatomical or radial-circumferential-longitudinal (RCZ) axes using an orthonormal transformation matrix (**Q**) as:

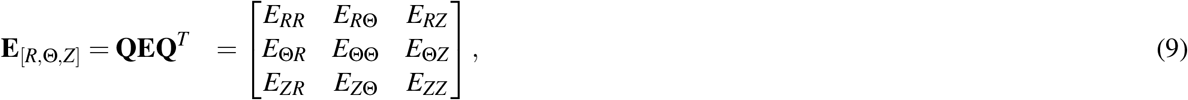

where *E*_*RR*_, *E*_ΘΘ_, and *E*_*ZZ*_ are the radial, circumferential, and longitudinal strains, respectively. Finally, the calculated strains were mapped onto a reconstructed geometry of the LV obtained through segmentation to facilitate the visualization of 4D strains. This visualization is achieved through Delaunay triangulation of all the pixels and strain quantities pertaining to the LV.

### 2.2 Application of SRR to the murine heart

#### 2.2.1 Image acquisition

Two distinct imaging protocols were devised to estimate 4D myocardial strains in C57BL/6 mice. The objective of the first protocol was to evaluate the effectiveness of the proposed SRR framework in improving myocardial strain reproducibility. In this protocol, imaging was performed in a three-month-old wild-type (WT) mouse (n = 1) and a three-month-old diabetic mouse (n = 1) [44]. Two different combinations of SA and LA scans were designed, with each set acquired one day apart. On the first day, five LA scans were acquired orthogonally, i.e., scans were acquired through translation between the anterior and inferior walls of the LV (Fig. 2A). The following day, five LA scans were acquired by radially sampling different image planes i.e., scans were acquired through rotating about the LV chamber (Fig. 2B). On both days, eight SA scans describing the region between the basal section of the LV and apex were acquired. The objective of the second protocol was to evaluate the effectiveness of the SRR framework in improving strain variability across a homogeneous cohort. For this protocol, we used a cohort of five six-month-old WT mice (n = 5). These mice were subjected to similar SA imaging, with LA scans acquired solely via orthogonal sampling. The mice used in both studies were males. In both protocols, cine CMR imaging was performed using a 7T Bruker Avance III HD scanner with a 72 mm quadrature proton volume coil (Bruker BioSpin MRI GmbH). In both protocols, an ECG signal was used to trigger image acquisition (R-R over three periods), and the combined SA-LA set was acquired in one session. The scan parameters were as follows: Fast Low Angle Shot, TE =1.6 ms, TR = 9 ms, flip angle = 20°, readout bandwidth = 65789 Hz, resolution = 200 µm, slice thickness = 1 mm, field of view = 32 x 32 mm, averages = 4. The imaging protocols were approved by the Cincinnati Children’s Hospital Medical Center Animal Care and Use Committee (Protocol 2018-0054).

**Fig. 2.**
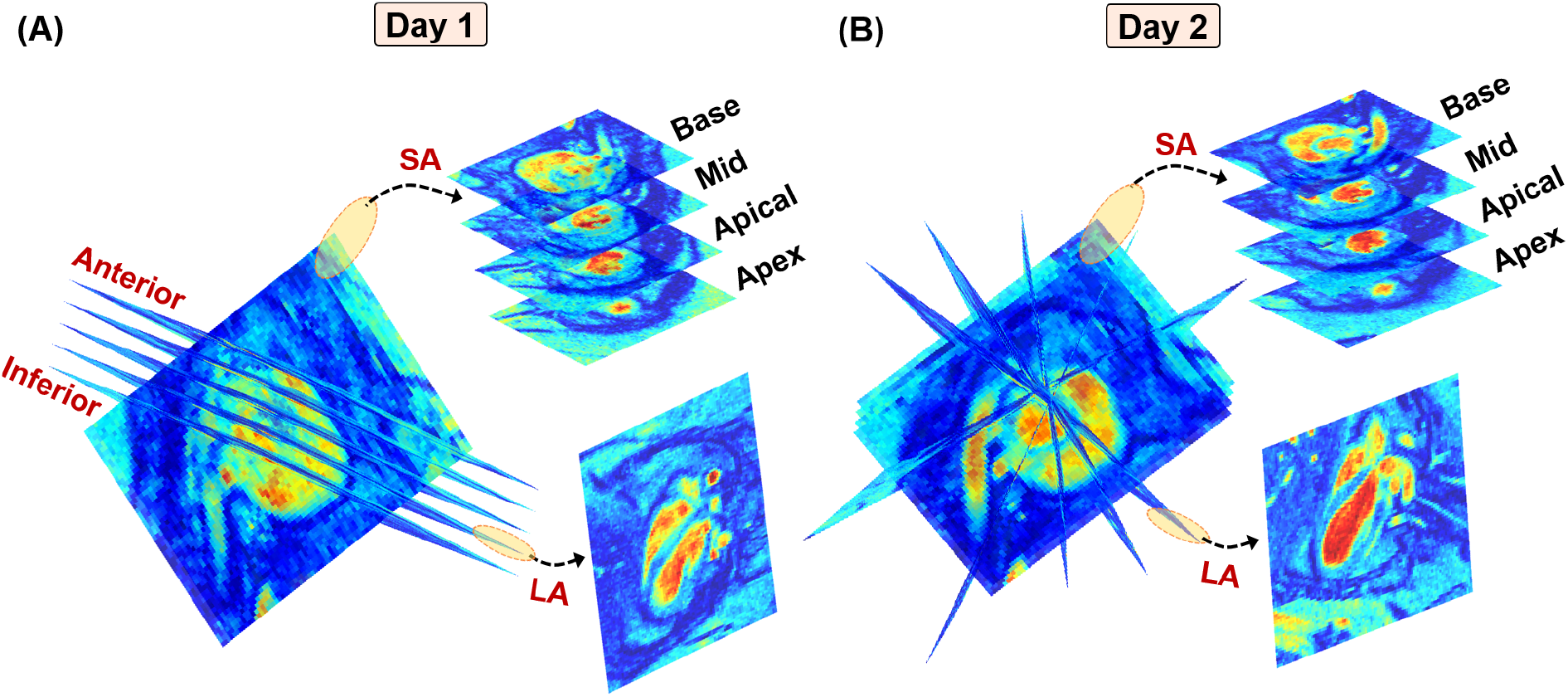
Imaging protocols used in the acquisition of cine CMR images for a three-month-old diabetic mouse over a period of two consecutive days. Combination of eight short-axis (SA) image planes between the basal slice and the apex of the left ventricle (LV) combined with five long-axis (LA) slices sampled (A) orthogonally between the anterior and inferior walls of the LV on the first day and (B) radially about the LV chamber on the next day.

#### 2.2.2 Myocardial strain estimation

Through the image acquisition protocols, various combined SA-LA sets were subjected to SRR, and subsequently, pixel-level displacements and strains were calculated at end-systole (ES), with end-diastole (ED) as the reference frame. Myocardial strains were evaluated separately for SR images, herein referred to as the SR model and the conventionally reconstructed LV. Here, conventional reconstruction refers to the use of image stacks that only contain LR SA images of the LV, herein referred to as the LR model. 2D cubic interpolation was used to increase the in-plane resolution of these LR images to match the in-plane resolution of the SR images. Each LR image was subjected to smoothing using a Gaussian kernel (*σ* = 2) before image registration. Unlike the SR images, longitudinal query points for the conventionally reconstructed images were determined strictly based on the aspect ratio of the SA scans [20]. Additionally, the SR grids resulting from the SA-orthogonal LA and SA-radial LA image stacks, herein referred to as SR-O and SR-R, respectively, were also used to calculate myocardial strains. In addition to mapping strains on murine-specific LV geometries, strains were also mapped in American Heart Association (AHA) standard segmentation maps, wherein the basal, mid, and apical sections of the LV were divided into 16 unique segments [45]. The AHA segmentation maps were used in calculating GLS and also investigating intra-cohort strain variability in strain distribution using LR and SR images of the WT mice.

### 2.3 Image quality analysis

Prior to strain estimation, the quality of the SR-O and SR-R images was assessed to evaluate the image reconstruction performance of SRR. Strain calculations have been observed to be sensitive to the quality of individual image planes, with artifacts such as discontinuities and blurring affecting the diffeomorphic demons algorithm. Specifically, the spatial definition (image sharpness) of the endocardial and epicardial borders in the reconstructed images was identified as a major contributor to strain reproducibility. Here, reproducibility refers to qualitative and quantitative similarities in strain patterns for the study on different days of image acquisition. The overall effects of each entity on strain estimation were grouped into three categories, (i) the endocardial sharpness, (ii) the epicardial sharpness, and (iii) artifact presence. Here, sharpness refers to the definition of the myocardial tissue at each region and was evaluated using image acutance, which is a subjective metric that relates the spatial resolution of an image to its spatial frequency. A small interrogation window describing the endocardium and epicardium was sampled, and acutance was evaluated using the image gradients such that:

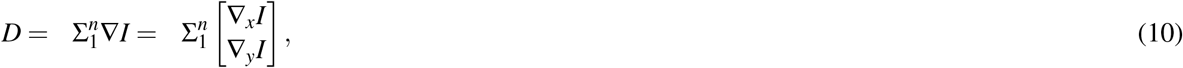

where **D** is the acutance, **I** is the interrogation window, and ∇_x_ and ∇_y_ are the gradient operators in the x and y directions of the window. Mean acutance was evaluated using the root-mean-squared (RMS) value of all the image gradients. A manual qualitative analysis in terms of a 5-point Likert scale (1 to 5) was performed for the SR images generated using the first imaging protocol, with each value signifying the levels of strain reproducibility. The lowest score denotes abject variability, and the highest indicates completely reproducible strains. The scoring scale for categories (i) and (ii) ranges between 1 = no distinct border due to poor acutance and 5 = perfectly defined border or high acutance. Scoring for artifact presence was determined between 1 = significant presence of unnatural features such as gradient discontinuities in the form of lines or the isolated presence of sharp pixels and 5 = no artifacts.

### 2.4 Validation using an in-silico phantom

The 4D strains generated through the SRR framework were validated using an *in silico* heart model phantom. Synthetic images of the heart were synthesized from 3D finite element (FE) simulations of mouse-specific cardiac contraction using the method described extensively in [46]. Briefly, the connectivity data of a meshed geometry was initially used to create an unstructured VTK grid in its undeformed state (end-diastole). Deformation at every subsequent timeframe was obtained by applying the FE-derived Cartesian displacement vectors onto each node, resulting in individual grids for each load increment. These grids were sectioned at incremental positions along the (a) z-axis to obtain SA slices and (b) its orthogonal axis to obtain LA slices and were projected onto a uniform grid of predefined parameters. The orthogonal axis was defined along the normal **n** = [1, 1, 0]. As a result, a stack of synthetic CMR images describing the deformation of the geometry was obtained. Given the predetermined number of load increments used in the FE solutions, each stack can be conceptualized as an individual temporal frame within an imaging sequence. Finally, the phantoms were exported as .tiff images using the lossless Lempel-Ziv-Welch compression algorithm with grid information (such as plane position, normal, projection, pixel spacing, and resolution) passed as metadata. Artificial SNR was added by convolving the grids with a Gaussian kernel. Thus, (i) a stack of SA images and (ii) a combined stack of SA and orthogonally sampled LA images were used to create two distinct sets of phantoms were created. While the first stack was used to obtain the myocardial strains conventionally, SRR was implemented using the second stack to evaluate the accuracy of LR and HR images in estimating myocardial strains.

## 3 Results

### 3.1 Reduction in through-plane anisotropy

The fundamental consideration of the SRR framework was to increase the spatial resolution of the CMR image stacks, with the benefits of SRR in through-plane LV reconstruction reported in this section. The reconstructed LV is shown across an arbitrary plane sampled perpendicular to the SA planes (Fig. 3). Since only eight LR SA slices were used in the image acquisition process, the LV reconstruction using LR images was hampered considerably by the slice thickness (Figs. 3A, B). Whereas, in both cases of SR images, the pixel spacing increased from 140 x 140 x 8 to (i) 140 x 140 x 110, for r_1_ = 1.0, and (ii) 280 x 280 x 220, for r_1_ = 0.5, respectively. Consequently, any number of planes could be sampled (Fig. 4) without being restricted to the orientation of the original LA scanning planes. As expected, there were no discernible differences between the LV chamber and the myocardial tissue along the through-plane direction of the LR images. In contrast, reconstruction was improved using SRR with a clear distinction between the high contrast blood pool and the relatively low contrast myocardial tissue (Figs. 3C-H) throughout the cardiac cycle (Fig. 4). However, when the in-plane resolution was limited to r_1_ = 1.0, a few artifacts were encountered in the reconstruction process (Fig. 3E, F). Despite a minor presence of artifacts, both SR protocols outperformed the LR image-derived LV reconstruction, irrespective of the resampling ratio or nature of the global interpolation scheme.

**Fig. 3.**
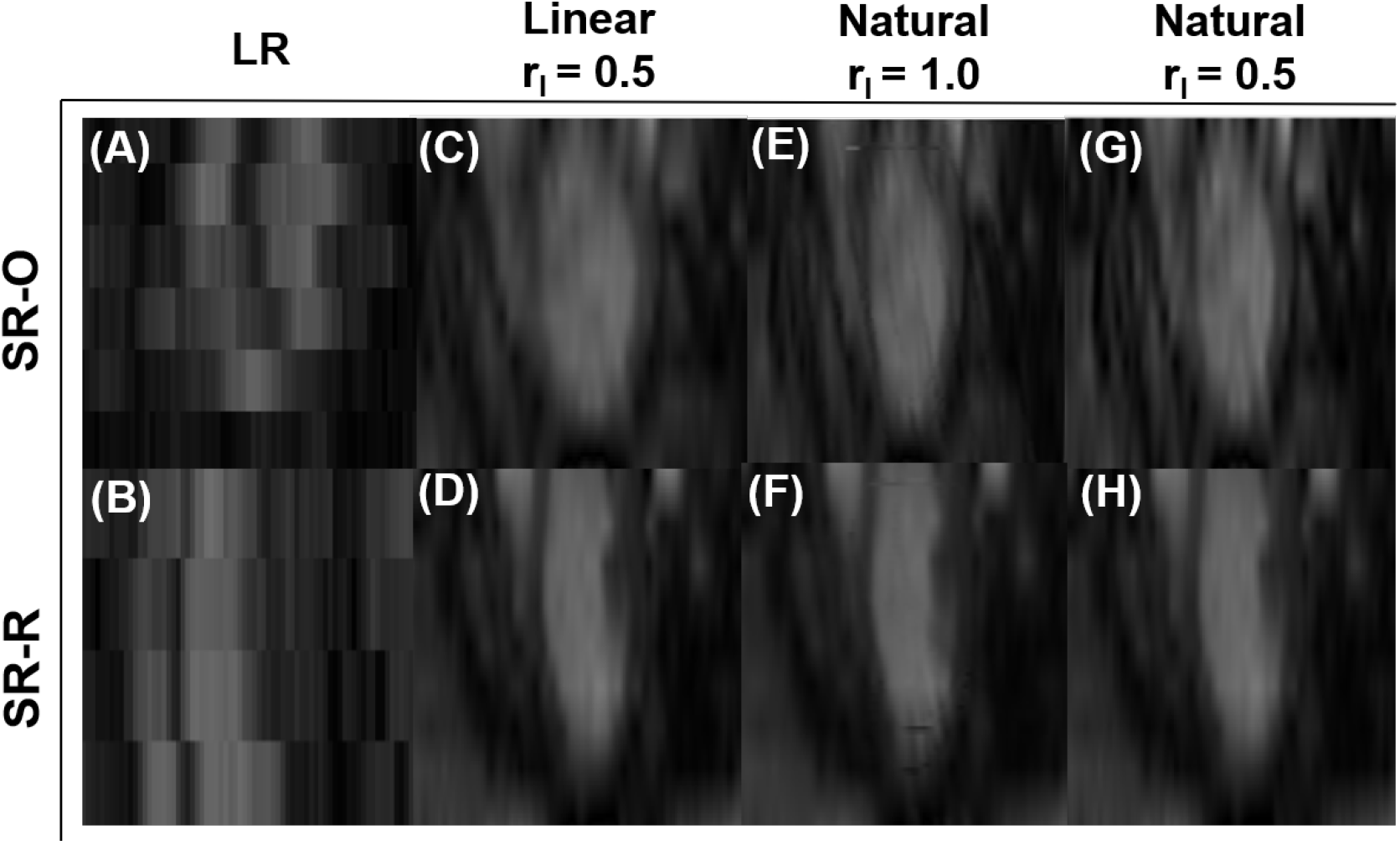
Through-plane reconstruction of an arbitrary plane perpendicular to the LV short-axis (SA). (A, B) Via stacking only the eight low-resolution (LR) SA image planes one over the other, (C-H) via SRR using (C, D) *linear* interpolation and (E-H) *natural* interpolation. SR-O: the combination of SA and orthogonally sampled LA images; SR-R: the combination of SA and radially sampled LA images. r_1_: resampling ratio.

**Fig. 4.**
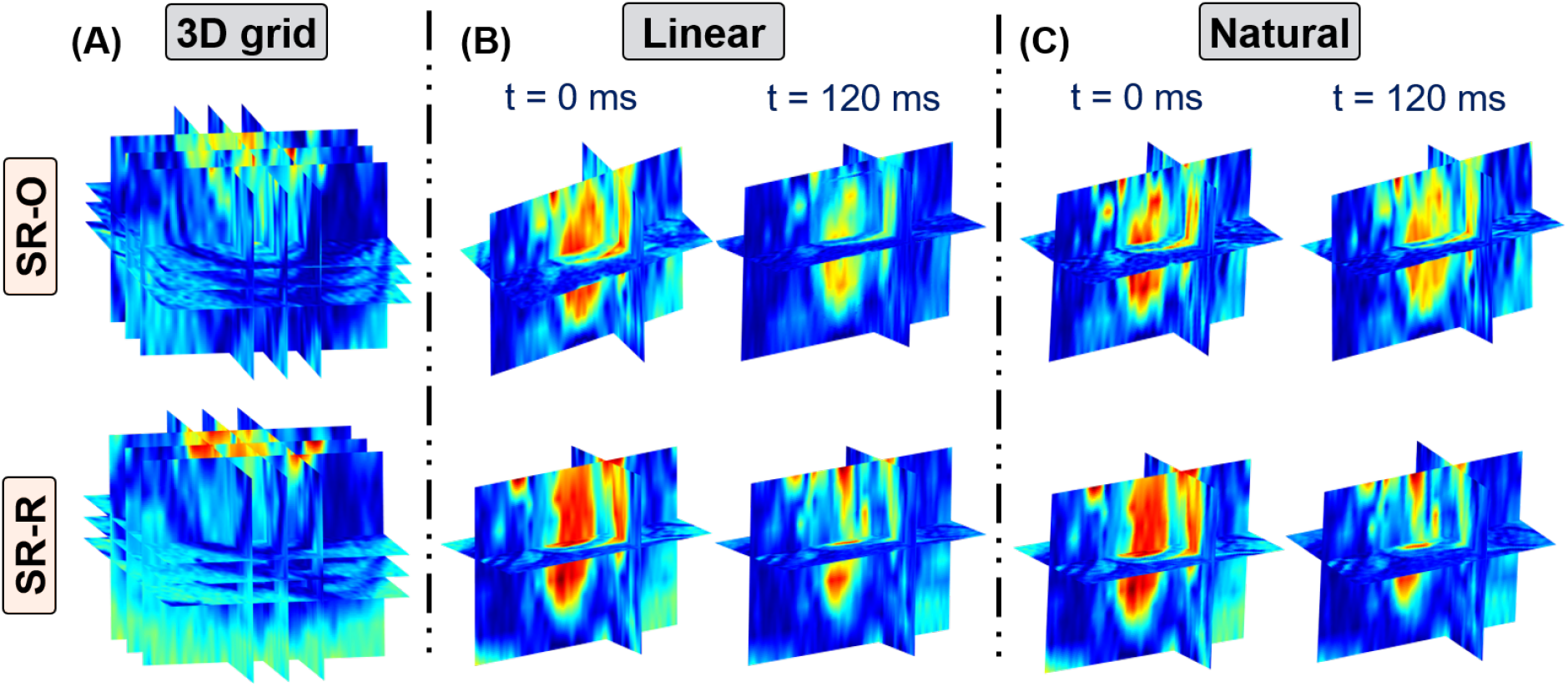
(A) Representative super-resolution reconstruction of the super-resolution (SR) image grid. The reconstructed LV is shown over arbitrary slices with the highest pixel intensities denoting blood. Representation of the cardiac cycle from (B) end-diastole (ED) at t = 0ms to (C) end-systole (ES) at t = 120ms over three orthogonal planes reconstructed using (top) SR-O and (bottom) SR-R. The SR grid was defined for a pixel spacing of 0.1 x 0.1 x 0.2 mm^3^ corresponding to a resampling ratio of r_1_ = 0.5. SR-O: the combination of SA and orthogonally sampled LA images; SR-R: the combination of SA and radially sampled LA images.

### 3.2 Improvement in tissue definition

#### 3.2.1 Qualitative assessment

The quality of SR-O and SR-R in reconstructing the eight original LR SA images was evaluated, and Likert scores were assigned based on the mean acutance values for both SR models generated using the *linear* and *natural* interpolation schemes. Results are also reported for their corresponding conventionally reconstructed models (Table 1). For both sets of SR models, the reconstruction of the epicardial and endocardial borders outperformed the LR models. Despite a pronounced improvement in the sharpness of the endocardial and epicardial borders, neither SR model was determined to have produced a perfect separation between the chamber and the myocardial tissue. This was expected as radial basis functions were used at the borders to regularize SNR-related errors. The most well-defined borders were produced through the *natural* interpolation scheme consistently, but this improvement in border definition was accompanied by a marked increase in artifacts, especially in SR-R (Figs. 5K,N-P). A few gradient discontinuities in the form of lines were observed over the SRR models. These lines were identified to be repercussions of isometric projections of the LR images onto the HR grid. These discontinuities were restricted to the original positioning of the LA arrangement. However, under the absence of in-plane resampling (r_1_ = 1.0), artifacts were observed over the entire image plane (Figs. 5C,K,O). The artifacts are reflected in the respective Likert scores, with the LR images outperforming SRR in this aspect as expected. Despite these artifacts, both SR models yielded overall improvements in both in-plane and through-plane reconstructions of the LV. However, qualitative comparisons for the LA reconstruction were not explored due to the insufficiency of LR models in delineating through-plane data (Figs. 3A and B).

**Table 1.** Qualitative analysis of SRR performance over a 5-point Likert scale. The original eight SA image planes were considered. All the results are presented as mean *±*SD. Linear and Natural denote the nature of the global interpolation scheme. SR-O: the combination of SA and orthogonally sampled LA images; SR-R: the combination of SA and radially sampled LA images.

| | | LR | Linear<br>($r_1 = 0.5$) | Natural<br>($r_1 = 1.0$) | Natural<br>($r_1 = 0.5$) |
| --- | --- | --- | --- | --- | --- |
| SR-R | Epicardium | $4.12 \pm 0.54$ | $4.20 \pm 0.21$ | $3.87 \pm 0.15$ | $4.43 \pm 0.23$ |
| | Endocardium | $3.73 \pm 0.31$ | $4.00 \pm 0.51$ | $3.53 \pm 0.56$ | $4.31 \pm 0.34$ |
| | Artifacts | $4.40 \pm 0.27$ | $4.14 \pm 0.24$ | $2.40 \pm 0.46$ | $3.97 \pm 0.38$ |
| SR-O | Epicardium | $4.32 \pm 0.18$ | $4.40 \pm 0.12$ | $3.88 \pm 0.21$ | $4.46 \pm 0.13$ |
| | Endocardium | $4.06 \pm 0.16$ | $4.11 \pm 0.14$ | $2.64 \pm 0.28$ | $4.45 \pm 0.12$ |
| | Artifacts | $4.6 \pm 0.18$ | $4.23 \pm 0.09$ | $2.51 \pm 0.17$ | $4.08 \pm 0.16$ |

**Fig. 5.**
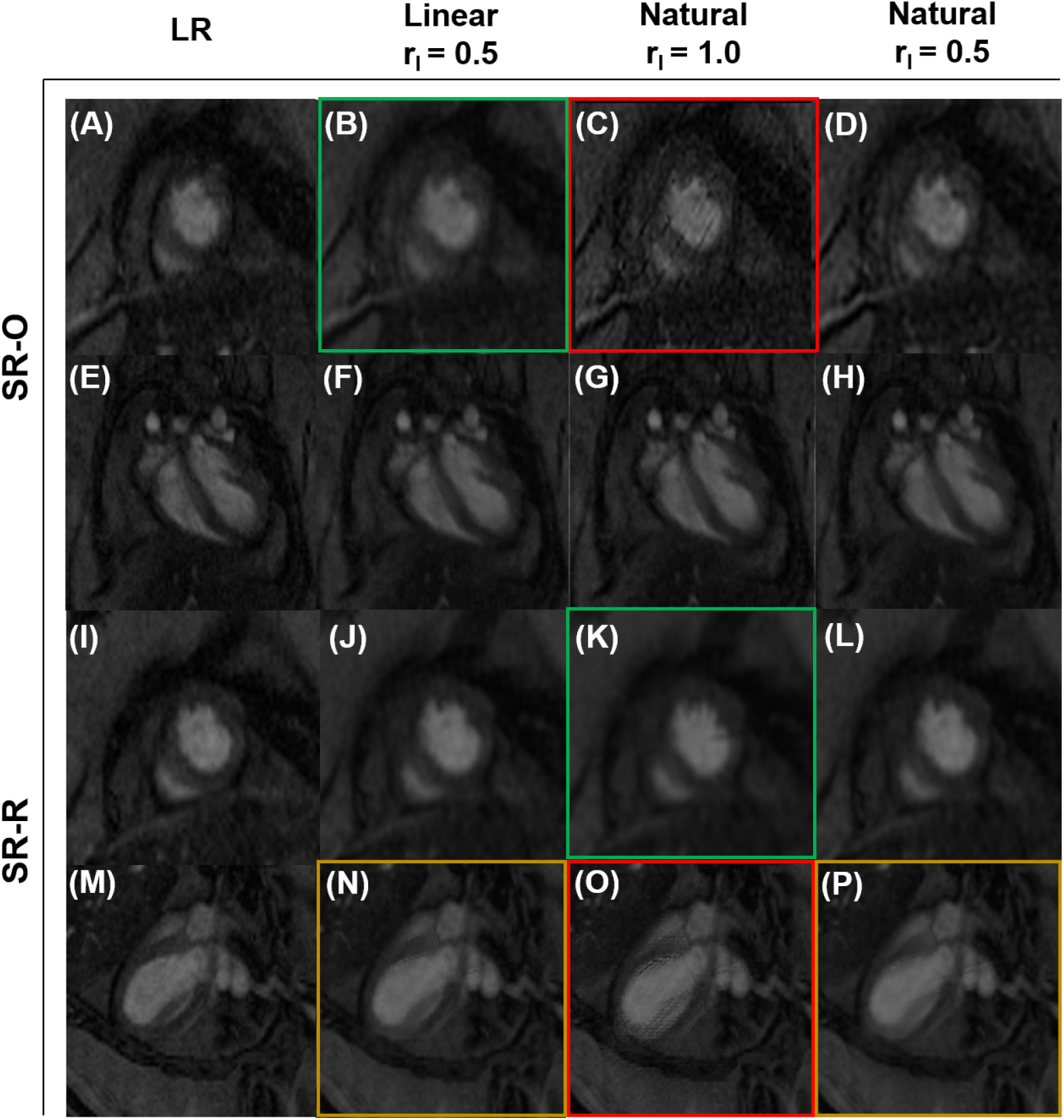
Super-resolution reconstruction of the original LR image planes of (A-D, I-L) the mid-SA slice, and (E-H, M-P) an arbitrary LA slice. Blurring and gradient discontinuities were observed with r_1_ = 1.0 contributing to the majority of the artifacts. Reconstructed image plane with the presence 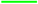 blurring, 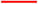 discontinuity artifacts, and 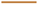 blurring & discontinuities. SR-O: the combination of SA and orthogonally sampled LA images; SR-R: the combination of SA and radially sampled LA images.

#### 3.2.2 Quantitative assessment

The quantitative significance of the artifacts on strain estimation was estimated through an analysis of the image quality metrics. The performance of SRR in reconstructing the images was determined using the structural similarity indices (SSIM) and root-mean-square errors (RMS). The metrics were calculated across all the SA and LA acquisitions for the SR models of r_1_ = 0.5, against the LR models (Table 2). SR images with r_1_ = 1 were compared to the original image acquisitions. Consistent with the qualitative assessment, *natural* interpolation models of SR with r_1_ = 0.5 produced images of high structural similarity with the original SA stacks (SR-R = 0.8526 *±*0.1172, and SR-O = 0.8234*±* 0.1187). Regardless of the interpolation schemes, SSIM of above 90% was encountered only at the SA images of the apical slice and the apex. Despite a relative absence of artifacts in the SR-O images (Fig. 5), higher levels of similarity were achieved using SR-R (Table 2). These SSIM values were likely to have stemmed from the similarity in blurring effects between the SR-R and LR images.

**Table 2.** Image quality metrics for SRR versus conventional reconstruction. The original LR SA (n_I_ = 8), and LA (n_I_ = 5) image planes were considered. The results are presented as mean *±*SD. Linear and Natural denote the nature of the global interpolation scheme. SSIM: structural similarity index, RMS: root-mean-square error. SR-O: the combination of SA and orthogonally sampled LA images; SR-R: the combination of SA and radially sampled LA images.

| | | Linear ($r_1 = 0.5$) | Natural ($r_1 = 1.0$) | Natural ($r_1 = 0.5$) |
| --- | --- | --- | --- | --- |
| SR-R | SSIM | $0.7856 \pm 0.1478$ | $0.715 \pm 0.1356$ | $0.8526 \pm 0.1172$ |
| | RMS | $0.1542 \pm 0.0841$ | $0.1851 \pm 0.1143$ | $0.1556 \pm 0.1087$ |
| SR-O | SSIM | $0.7743 \pm 0.1345$ | $0.7045 \pm 0.1085$ | $0.8234 \pm 0.1006$ |
| | RMS | $0.1523 \pm 0.0846$ | $0.2098 \pm 0.1119$ | $0.1234 \pm 0.1187$ |

### 3.3 4D myocardial strain distribution in the murine heart

#### 3.3.1 SRR suppresses unrealistic strain estimation

4D strains were calculated using the SR and LR models at various time points describing the cardiac cycle, and a summary of arbitrary SA and LA images describing the cardiac cycle is presented in the supplementary material (Supplementary Figs. S1 and S2). The resulting strain maps are reported for both the diabetic (Figs. 6-8) and WT mice (Supplementary Fig. S3). In contrast to the conventional LV models, transmural and regional variations in strains were quantified over the entire LV using SRR (Figs. 6-8). While strains have been reported for both models, strains for the conventionally reconstructed LV (Figs. 7A-C, G-I and 8A-C, G-I) were merely mapped onto the SR models using a nearest neighbor approximation. A visualization of the epicardial strains has been presented using only the SR models, as the LR models revealed unrealistic through-plane strains. As per literature [47], any strain over a limit of 50% was designated to be non-physical. At ES, regional epicardial strains using SR showed varying degrees of contraction between the base and the apex, with apical shortening exceeding the other regions (Fig. 6). Clinically relevant estimates of strains are often measured at the basal, mid, and apical slices. Accordingly, the circumferential (Fig. 7) and longitudinal (Fig. 8) strains were visualized at these slices and along the endocardium. Although qualitative similarity in strain distribution was observed between the two methodologies, there was a significant difference in the peak strains. Specifically, for the LR model, there was a significant presence of positive longitudinal strains despite global LV shortening (Septal; LR, E_ZZ_ = 0.7342 vs SR-O, E_ZZ_ = 0.4985. Lateral; LR, E_ZZ_ = 0.5146 vs SR-O E_ZZ_ = 0.3126). Also, the regional circumferential shortening seen in SR was recapitulated in the LR model with similar differences in peaks. However, there was an abundance of non-physical transmural strains, which were suppressed by both SR-O and SR-R models (Figs. 7C, F).

**Fig. 6.**
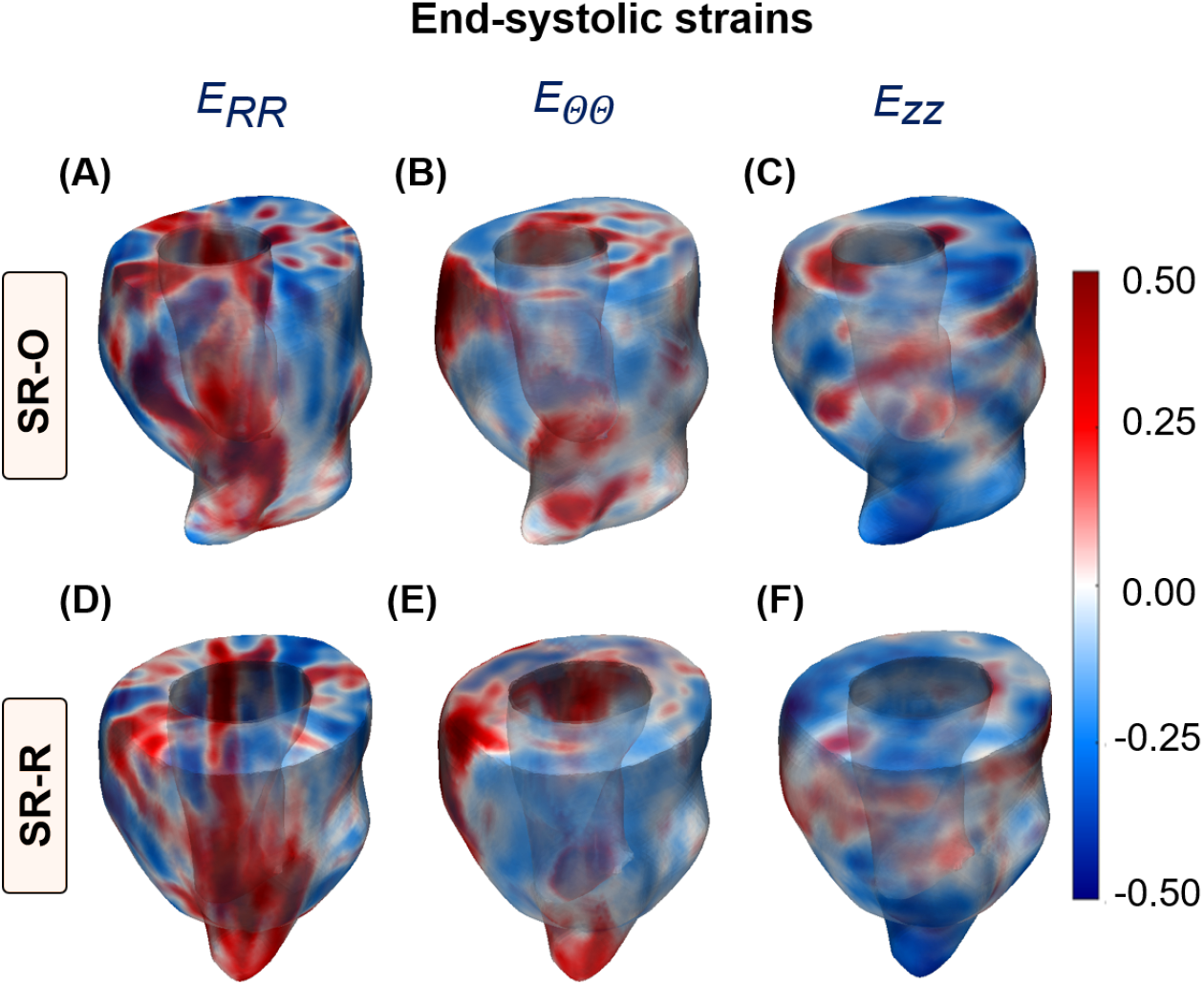
Myocardial strains along the radial (*E*_*RR*_), circumferential (*E*_ΘΘ_) and longitudinal (*E*_*ZZ*_) directions of the LV for (top) SR-O, and (bottom) SR-R at ES. Strains were mapped onto a reconstructed LV with a significant presence of positive radial deformation accompanied by regional distribution of negative strains in the circumferential and longitudinal directions in both reconstructions. SR-O: the combination of SA and orthogonally sampled LA images; SR-R: the combination of SA and radially sampled LA images. LR: low-resolution, SR: super-resolution

**Fig. 7.**
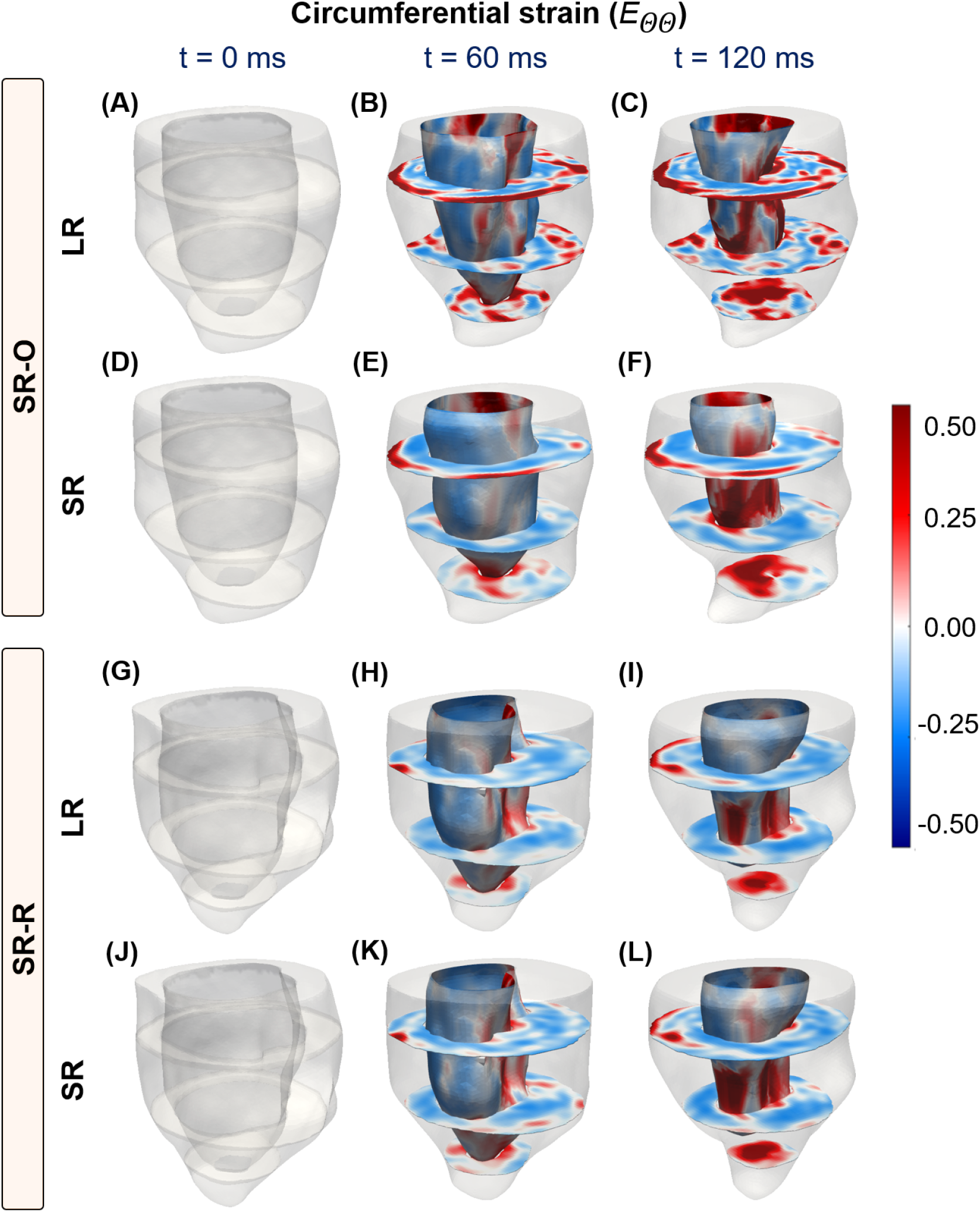
Time-course progression of circumferential strains from ED (t = 0 ms) to ES (t = 120 ms) at the basal, mid, and apical short-axis (SA) slices, and the endocardial wall of the LV for (A-F) SR-O, and (G-L) SR-R-generated models. (A-C, G-I) Reconstruction using the stack of LR SA images. (D-F, J-L) Super-resolution reconstruction. SR-O: the combination of SA and orthogonally sampled LA images; SR-R: the combination of SA and radially sampled LA images. LR: low-resolution, SR: super-resolution

**Fig. 8.**
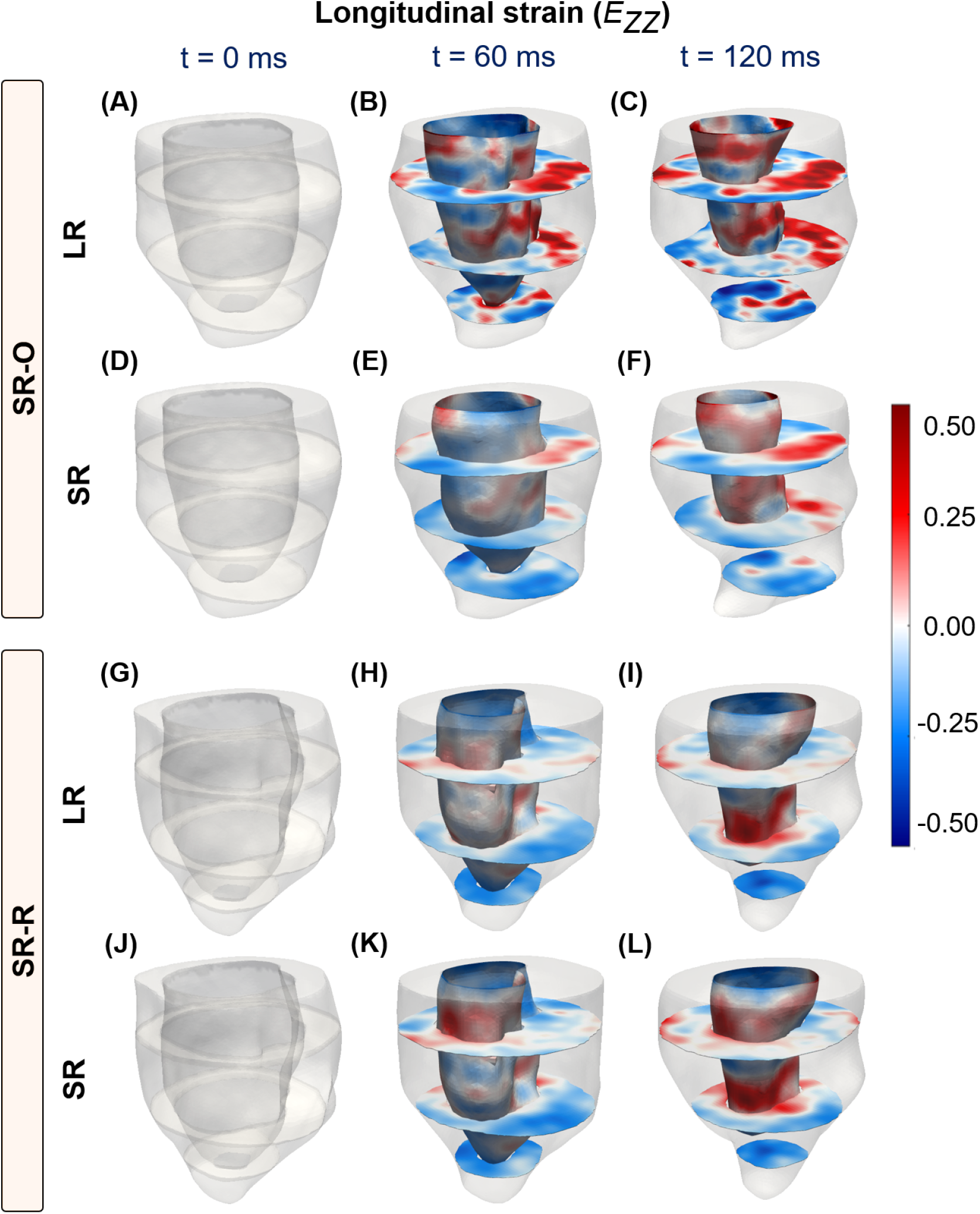
Time-course progression of longitudinal strains from ED (t = 0 ms) to ES (t = 120 ms) at the basal, mid, and apical short-axis (SA) slices, and the endocardial wall of the LV for (A-F) SR-O-, and (G-L) SR-R-generated models. (A-C, G-I) Reconstruction using the stack of LR SA images. (D-F, J-L) Super-resolution reconstruction. SR-O: the combination of SA and orthogonally sampled LA images; SR-R: the combination of SA and radially sampled LA images. LR: low-resolution, SR: super-resolution

#### 3.3.2 SRR improves the spatial and intra-cohort homogeneity of myocardial strains

The time-course homogeneity in regional myocardial strain distribution estimated between the SR-O and SR-R images and corresponding LR images are reported. As opposed to the considerable variations in strain patterns that were noted between the two LR models (Figs. 7C, I and 8C, I), the SR images provided more consistent strain distribution between both acquisition strategies. Variations in the regional contractile patterns were observed between the SR and LR models. For instance, large areas of circumferential thickening, concentrated in the lateral regions, were removed in the SR-O models, with the mean strain in all three sections of the LV reducing considerably (Lateral; LR, *E*_*θθ*_ = *−*0.0870 *±* 0.0140 vs. SR, *E*_*θθ*_ = *−*0.1541 *±*0.0277). There was a great agreement in the contractile patterns between SR-O and SR-R, with the lateral regions of the LV showing similar contractile patterns indicating improvements in reproducibility (Lateral; SR-O, *E*_*θθ*_ = *−*0.0870 *±* 0.0140 vs. SR-R, *E*_*θθ*_ = *−*0.07124 *±*0.0148).Moreover, intra-cohort strain variability investigated as the degree of spatial heterogeneity in the regional strain distribution is reported for five WT mice (Fig. 9). The suppression in unrealistic strain patterns was observed across all mice through the SR models (Fig. 9A), and estimations of GLS for conventional (LR) strain calculations tended to show higher variability across five mice compared to that of SR measurements (Fig. 9B). Similarly, SR demonstrated considerable improvements in minimizing the standard deviation (SD) at each segment between the mice (Fig. 9C), with a 50% reduction in the coefficient of variation measured as the ratio of the SD to the mean at each segment (Fig. 9D; SR vs LR: −0.2186 vs. −0.5732).

**Fig. 9.**
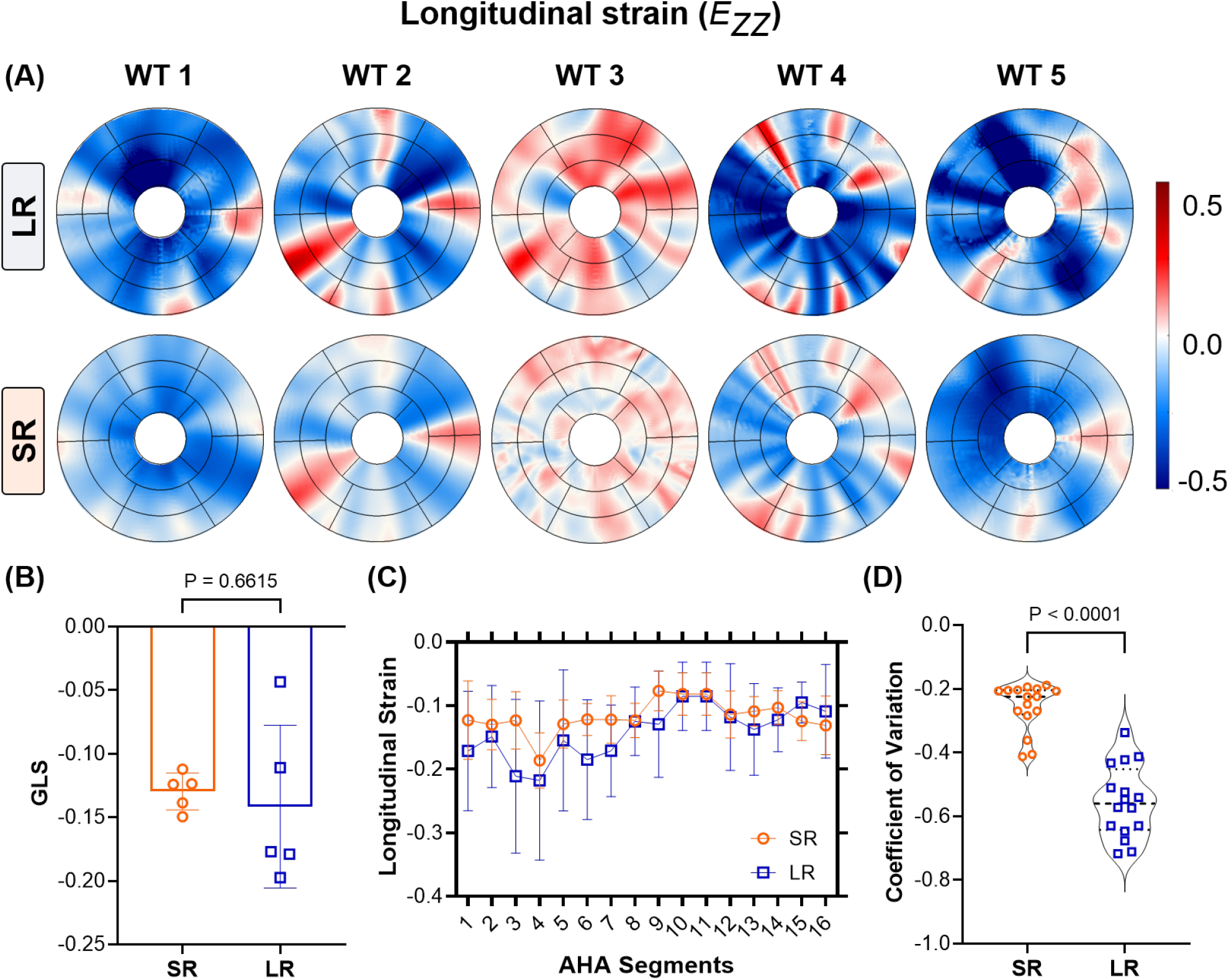
Evaluation of the spatial homogeneity of myocardial strain distribution using low- and super-resolution imaging in wild-type mice (n = 5). (A) Regional distribution of end-systolic longitudinal strain, presented in AHA segmentation plots at the base, mid, and apical slices. (B) Estimation of global longitudinal strain (GLS) as the average of all regional strain quantities in each AHA segment. (C) Comparison of the average longitudinal strain in each AHA segment, with the values represented as mean *±*standard deviation (SD). (D) Violin plots of the coefficient of variation (CV) of the strain distribution at each segment for all mice. LR: low-resolution, SR: super-resolution, WT: wild-type.

### 3.4 Validation of SRR in CMR using an in silico phantom

The accuracy of the SRR framework in calculating myocardial strains was validated using an SR phantom generated using a stack of eight SA images and five orthogonally sampled LA images and compared to a stack of LR SA images (10A and B). The accuracy of the respective stacks was estimated using mean-squared error (MSE) analyses, and readings are reported for the time-course progression of contractile strains described by both phantoms at the apical, mid, and basal planes (Fig. 10 C-H). Both phantoms presented comparable errors in the estimation of end-systolic *E*_*RR*_ (LR vs SR = 9.4033 vs 10.0104). However, improvements were noted in the calculations of both *E*_*θθ*_ (LR vs SR = 12.5700 vs 9.9553) and *E*_*ZZ*_ (LR vs SR = 26.2933 vs 15.0667) at ES. Additionally, improvements due to SRR were observed in the early stages of contraction for all three strain quantities.

**Fig. 10.**
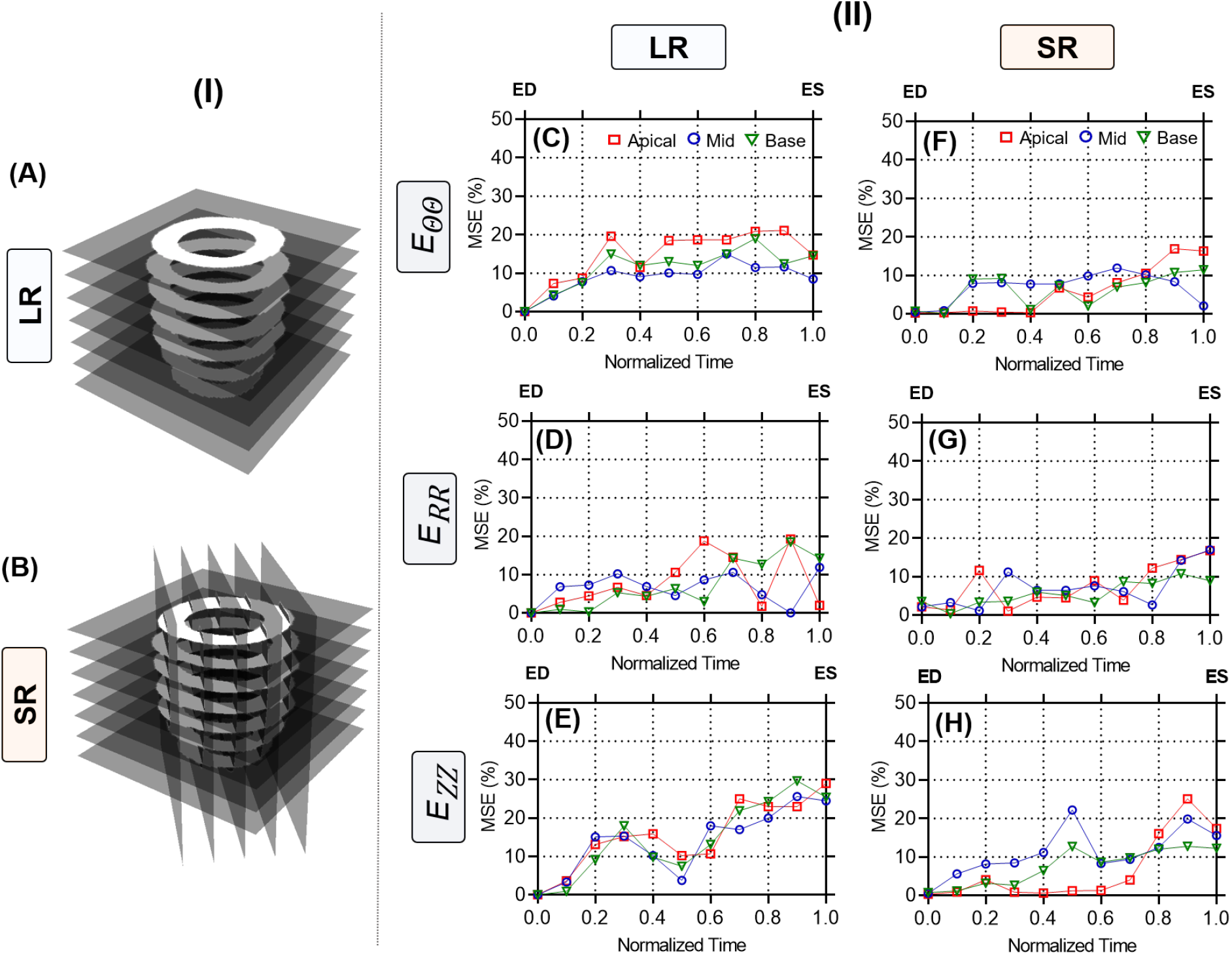
Mean squared error (MSE) curves comparing the errors in strain estimation between the image-derived and ground-truth strains established via an in-silico phantom at various timepoints between end-diastole (ED) and end-systole (ES). Strains were calculated through image registration using (A) low-resolution (LR) and (B) super-resolution (SR) phantom images. MSE curves for all three strain quantities are presented at three sectional planes of the LV derived using the (C-E) LR and (F-H) SR phantoms. *E*_*θθ*_ : circumferential, *E*_*RR*_ : radial, and *E*_*ZZ*_ : longitudinal strains.

## 4 Discussion

### 4.1 Establishing 4D regional LV strain maps

CMR-based methods of strain estimation in the LV are influenced primarily by substandard delineation of in-plane cardiac motion and insufficient data for through-plane measurements. These limitations have contributed to facile 2D in-plane representations of the complex LV motion. While some strategies have extrapolated longitudinal deformation by interpreting deformation patterns at the endocardial and epicardial layers [19], others have used contrast-enhanced image acquisitions to assess regional distributions of 3D strains in SA and LA images [48]. However, there still remains a paucity of literature on the quantification of 4D myocardial strains. In this study, the structural characterization was sought to aid in developing a complete 4D LV structure-function relationship. SRR-based methods have been validated in their abilities to improve image quality and have recently been studied for their capabilities in reducing CMR acquisition times [34] and improving diagnostic reliability [35]. However, the applications of SRR in assessing LV deformation remain vastly under-explored. We presented a high-fidelity pipeline to characterize the 4D regionality of LV function by leveraging combined acquisitions of SA and LA images with the SRR framework and investigated the effects of SRR on strain calculations. A priori knowledge in the form of tissue class was used to dissect the classical SRR problem and enhance the reconstruction of the LV geometry. Comparisons with conventional LR images of the LV that exhibit significant through-plane anisotropy revealed marked differences in the estimation of in-plane and through-plane strains (Figs. 7, and 8). Strain quantities estimated via the LR and SR models were validated by using an in-silico phantom, and SRR was noted to improve the description of through-plane motion. Additionally, Our findings showed the recapitulation of expected global characteristics of LV shortening, an overall reduction in unrealistic strain values, improvements in the intra-cohort homogeneity of myocardial strains, and a significant reduction in through-plane anisotropy, with minimal degradation of tissue contrast.

### 4.2 Improving through-plane strain accuracy using enhanced image acquisition

The LV exhibits twisting during systolic contraction, resulting in significant through-plane deformation with corresponding in-plane mechanical adjustments via radial thickening and circumferential shortening. These deformations translate as “out-of-plane” pixel motions that are not captured by traditional CMR imaging of the LV. Standard functional imaging is often limited to 7-9 SA planes with an optional LA or four-chamber view. The resulting pixel set is heavily anisotropic, which impediments accurate strain calculations. We hypothesized that an SRR-based methodology combining scan planes of varying orientations would considerably reduce pixel spacing, thereby improving strain calculations. In the evaluation of longitudinal strains, a significant presence of positive regional strains was noted in the LR model (Figs. 8 and S3) despite similar global shortening (Fig. 9). Notably, the LV was shown to elongate at ES when strains were generated using the SA stack (Figs. 8C, I). These observations were expected as the through-plane anisotropy distorts displacement derivatives and subsequent strains (Eq. 8). The interpolation module of the SRR framework was specifically designed to address this concern. Piecewise cubic interpolants have been shown to be effective in reconstructing 2D image planes from non-uniform datasets [49, 50]. The *natural* or *pchip* interpolation method showed very minute RMS errors and high SSIM (Table 2) in reconstructing individual image planes. Despite significant upsampling of query points in all directions, average SSIM values of 80% were still achieved with very minimal artifacts (Table 1). Building on these minor deviations in feature preservation, the pixel spacing using SRR was increased to about 25 times the original acquisition. Through this nearly isotropic pixel set, apical and endocardial shortening was captured more accurately in both SRR models at ES (Figs. 8F,L), with a very minor presence of positive longitudinal strains across all WT mice (Fig. 9). The benefits of SRR in improving longitudinal shortening were validated using the in-silico phantom while minimizing errors at three different planes of the LV. These errors can be attributed to motion artifacts, especially at the tissue-pool interface of the endocardium, that confound image registration. Despite the presence of these artifacts, errors in all strain quantities were restricted to under 10%, highlighting the potential improvements in 4D strain calculations.

### 4.3 The effects of edge sharpness on in-plane motion calculation

The standardization of regional strain calculations using any imaging modality is challenged by inter-study and inter-vendor variability. These limitations are either caused by assumptions in delineating cardiac motion, which differ between calculation strategies, or, in most cases, are dependent on the operator’s capabilities. The inadequacy of temporal resolution in standard CMR images results in sharp gradient changes between time points, thereby producing errors in registering consecutive image frames. A strong correlation was noted between sharp image gradients and the generation of non-physical strains, which necessitate regularization. A consequence of regularizing the image registration process was the underestimation of strains, especially at the tissue-pool interface of the endocardium. In the absence of this regularization, notable variations were observed between the two LR models of 4D strains with pronounced differences in both the peak values and the patterns (Figs. 7C, I, and 8C, I). The images on the second day (SR-R) were noticeably blurred compared to the first (Figs. 5A, I), resulting in a disparity in endocardial sharpness and subsequent strain estimation. By defining a convex hull of pixels with boundaries at the endocardium, the edge sharpness was increased using SRR, and the chamber pixels were isolated from the myocardial tissue. The diffusion of gradients across the edges was thus inhibited, and the effects were translated as such in the strain calculations (Equations **??**, 8). Given the minimal presence of artifacts in the SR-O images, the protocol was implemented in the study homogeneity of 4D strains in a cohort of WT mice. As opposed to the significant deviation in strains derived from the LR model, similar contraction patterns were observed between the different WT mice using the SR images (Fig, 9), indicating a pathway toward the reproducibility of 4D regional strains. However, contrary to the improvements in transmural strain accuracy, calculations at the endocardial and epicardial borders can still be improved since the corresponding pixels were subjected to regularization through radial basis functions to maintain continuity in the global interpolation scheme at the tissue-pool interface.

### 4.4 Potential of 4D strains for prognostic assessment of the myocardium

Heart diseases such as myocardial infarction and hypertrophic cardiomyopathy leading to heart failure are clinically diagnosed through global and regional volumetric and functional indices. Measurements of diastolic and systolic LV elastances, as well as diffusion tensor imaging and histological analyses, have confirmed mechanical and architectural changes in the LV due to a variety of cardiac remodeling mechanisms. These myocardial adaptations lead to alterations in cardiac motion and can be detected by quantifying myocardial strains. In cases of radiation therapy-induced cardiotoxicity, functional impairments are not immediately apparent through EF measurements. In the absence of worsening EF, regional strain calculations have been shown to be sensitive markers, especially at an early stage [51]. However, the innate complexity of regional cardiac motion, resulting from intricate myocardial architecture and passive and active phases of motion [52], remains to be a prominent challenge in rigorously quantifying full 4D cardiac motion in vivo. Image-based strain models usually restricted to 2D planes often yield confounded representations of the resulting myocardial deformation. The complex nature of 4D ventricular kinematics [53] demands the establishment of a standardized quantification methodology. The regional strain analysis presented herein carries the potential to establish a reproducible 4D representation of myocardial deformation. Through the SRR framework presented in this study, significant improvements in image quality, and strain calculations were achieved. This rigorous quantification of 4D strains can further be used in conjunction with real-time volumetric, and hemodynamic readings to estimate the intrinsic material properties of the myocardium [54]. Thus, a multiscale mathematical approach comprising organ-level measurements, and tissue-level mechanical characteristics can be posed to assist in clinical intervention.

### 4.5 Limitations

Despite the proposed utility of the SRR framework in promoting the prognostic efficacy of 4D myocardial strains, there exist a few limitations. First, investigations into the intra-cohort variability of 4D contractile patterns were restricted to WT mice. While this allowed us to present comprehensive image reconstruction and 4D strain reproducibility within controlled conditions, the implementation of the SRR framework in additional subjects is surely needed to further evaluate its performance and optimize its application across different pathophysiological conditions. Second, we were limited by the imaging capabilities in acquiring HR scans as a ground-truth metric to evaluate the performance of SRR in image reconstruction. However, this is common in most SRR methods [32, 35, 49] as capturing these acquisitions is limited by factors such as hardware capabilities (e.g, limited scan speeds) and government regulations. Finally, the SRR framework was susceptible to the generation of artifacts during image-plane reconstruction. Currently, the SRR framework relies on standard scattered data interpolation to generate HR images. Since the HR images were generated through interpolation, we noted artifacts in the form of discontinuous straight lines across the SA slices (Fig. 5). Crucially, we suspect these artifacts to have influenced the derivation of the image gradients and, subsequently, the strain estimation. We expect these artifacts to be reduced by implementing deterministic interpolation schemes [55, 56], yet to be implemented in MRI scans, or by using physics-informed image registration algorithms [57]. Alternatively, individual slices could be subjected to edge-enhancing diffusion filters. These filters have been known to reduce artifacts in image reconstruction by impeding diffusion across edges and have also been implemented in MRI scans of the brain [58, 59].

## 5 Conclusion

In this study, we presented a novel and comprehensive analysis of 4D myocardial deformation through the CSRR framework. The framework offers a translational tool that leverages combined CMR acquisitions and tissue classification strategies to improve LV reconstruction with minimal noise and artifacts. A detailed representation of 4D myocardial strains was made possible with such analysis, offering the potential to establish a continuous 4D LV structure-function relationship as a standard practice. To the best of our knowledge, this is the first application of any SRR methodology to enhance the in-vivo characterization of soft-tissue mechanics.

## 6 Acknowledgements

R. Avazmohammadi was supported by the NHLBI grant R00HL138288. J. Ohayon was partially supported by the Agence Nationale de la Recherche (ANR), through the SIMR project (ANR-19-CE45-0020). T. Mukherjee was supported by the American Heart Association (AHA) predoctoral fellowship 24PRE1240097.

## 8 Data availability

The data that support the findings of this study are available from the corresponding author, R.A., upon request.

## 9 Supplementary material

**Supplementary Figure S1.**
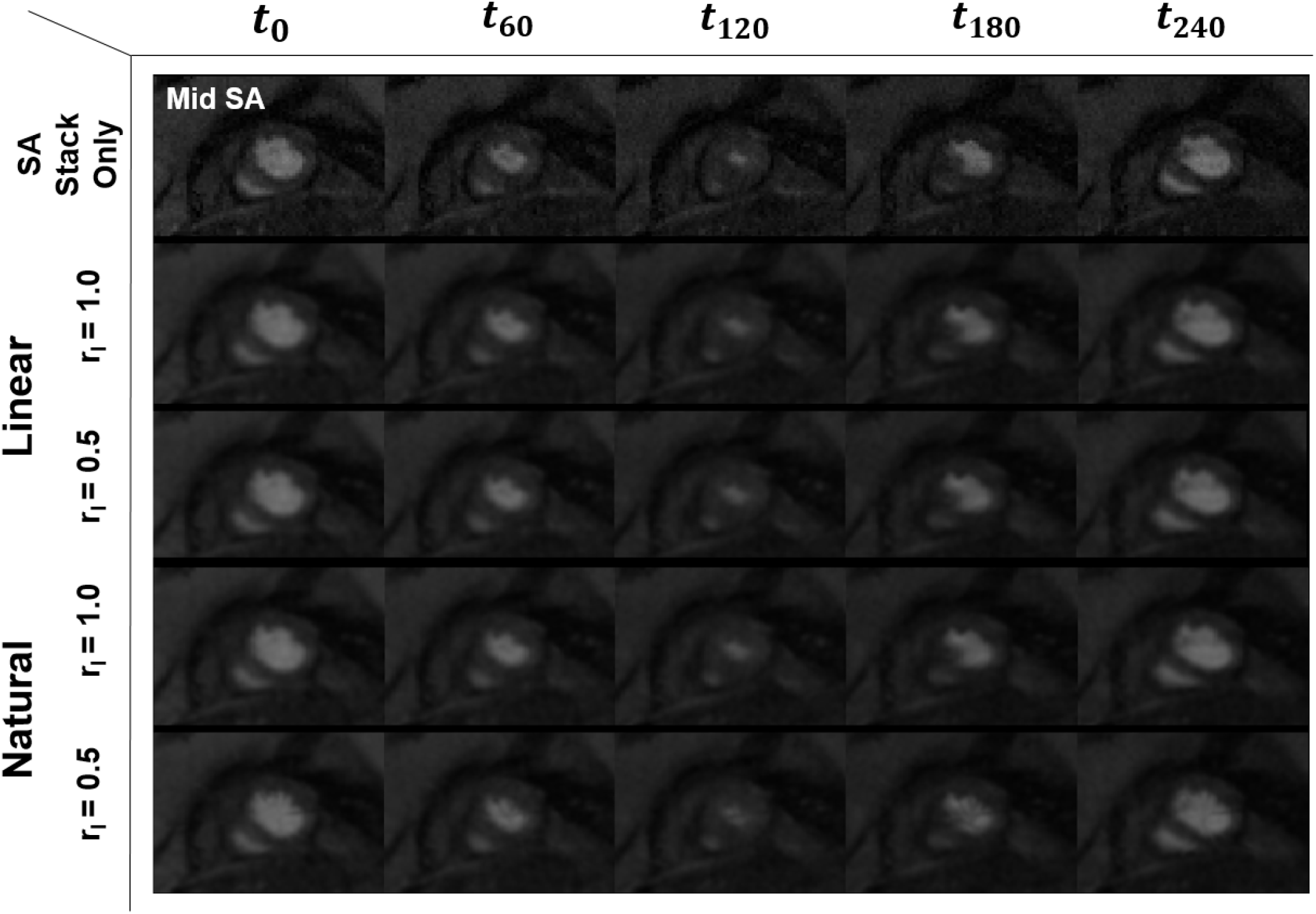
Super-resolution reconstruction of the mid-short-axis plane of a murine heart using the combination of SA and orthogonally sampled LA images. Images are shown for select timepoints within a cardiac cycle from ED to ES to ED with the total duration of t = 240ms. SA stack only represents the original low-resolution acquisitions, with Linear and Natural denoting the nature of the global interpolation. SR-O: the combination of SA and orthogonally sampled LA images; r_1_: resampling ratio.

**Supplementary Figure S2.**
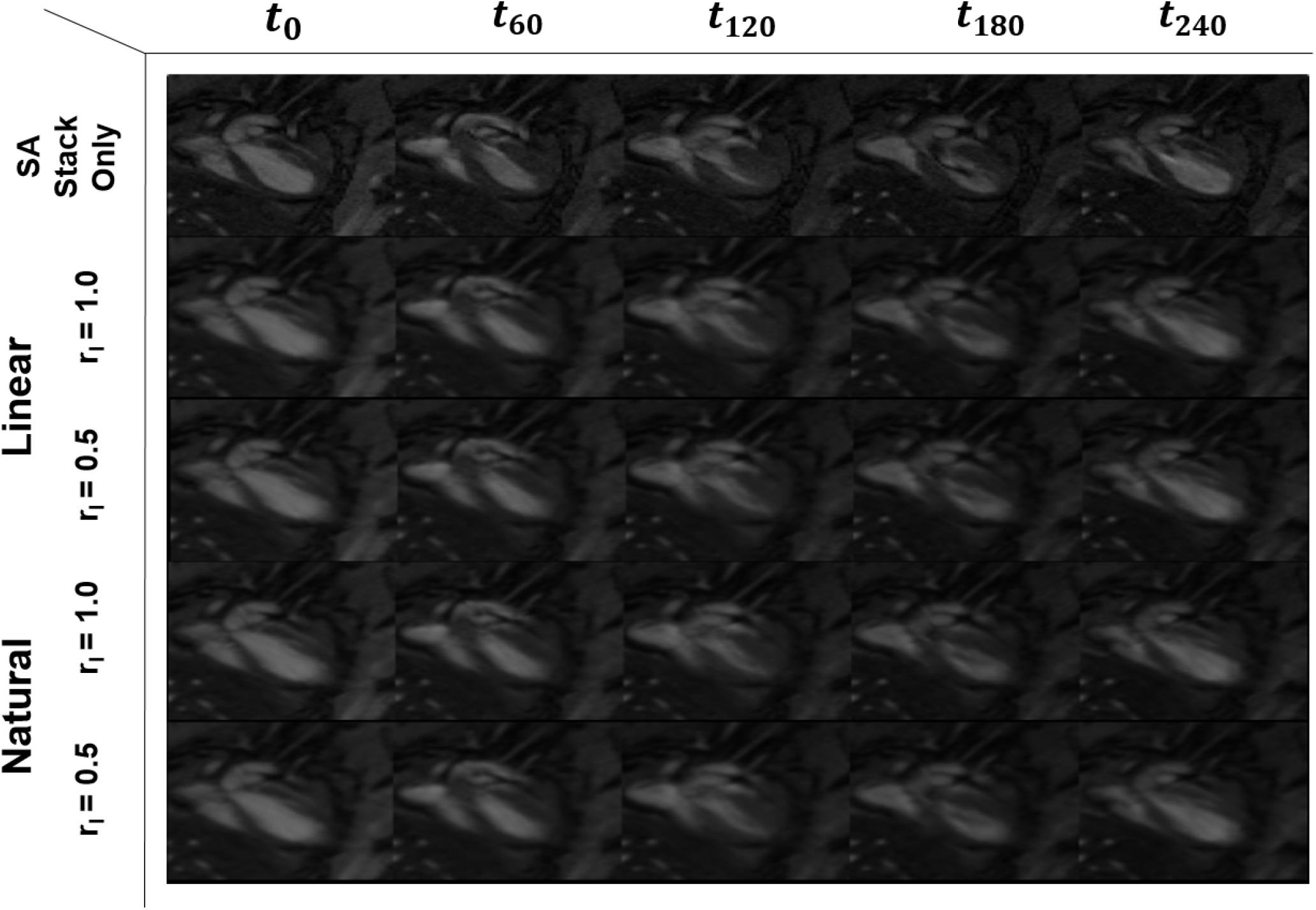
Super-resolution reconstruction of an arbitrary long-axis plane of a murine heart using the combination of SA and orthogonally sampled LA images. Images are shown for select timepoints within a cardiac cycle from ED to ES to ED with the total duration of t = 240ms. SA stack only represents the original low-resolution acquisitions, with Linear and Natural denoting the nature of the global interpolation. SR-O: the combination of SA and orthogonally sampled LA images; r_1_: resampling ratio.

**Supplementary Figure S3.**
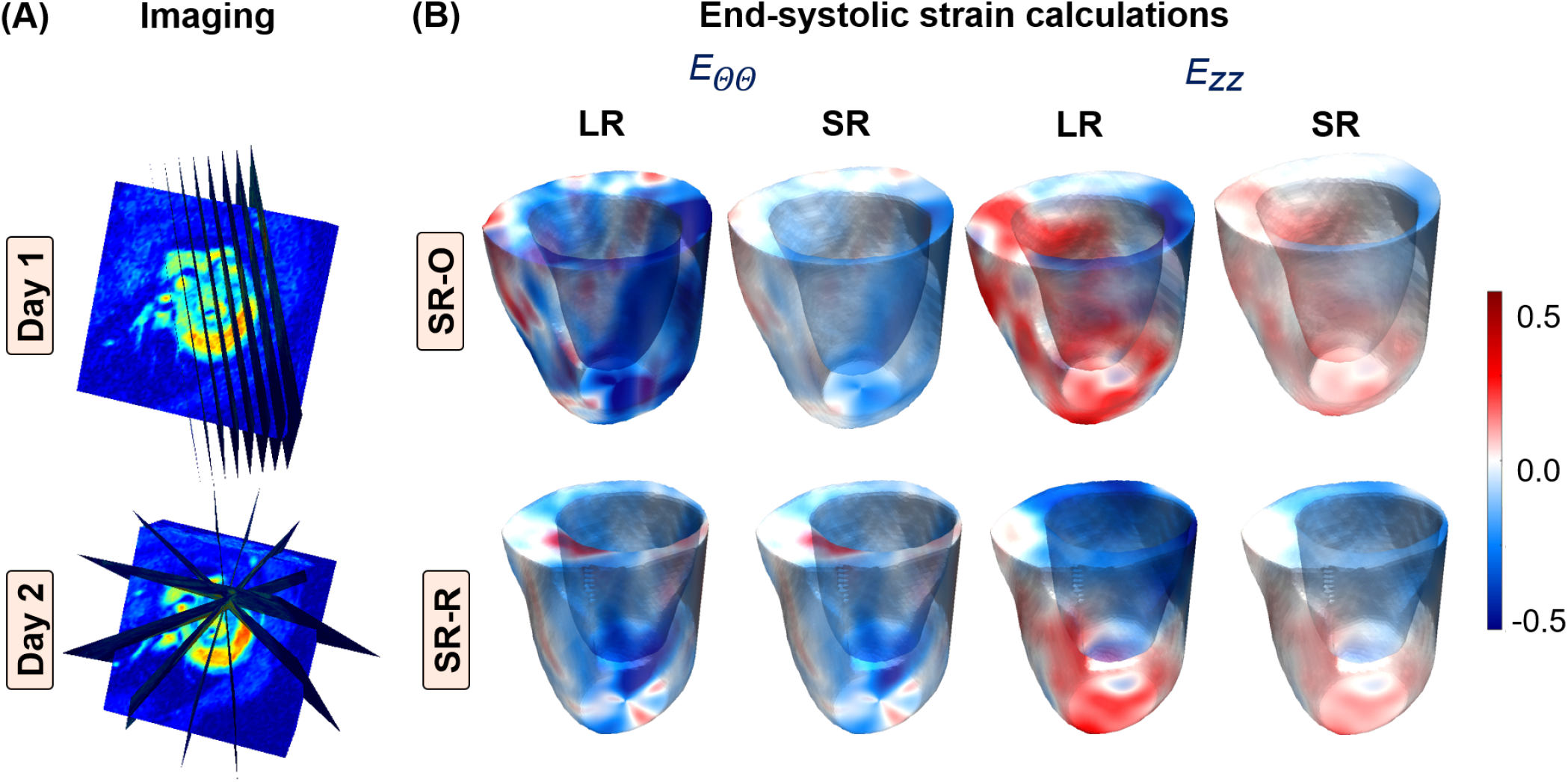
(A) Imaging protocols used in the acquisition of cine-CMR images for a three-month-old wild-type (WT) mouse over a period of two consecutive days. Combination of eight short-axis (SA) image planes between the basal slice and the apex of the left ventricle (LV) combined with five long-axis (LA) slices sampled (top) orthogonally between the anterior and inferior walls of the LV on the first day and (bottom) radially about the LV chamber on the next day. (B) Myocardial strains calculated for the same WT mouse along the circumferential (*E*_ΘΘ_) and longitudinal (*E*_*ZZ*_) directions of the LV from CMR images acquired on (top) Day 1 and (bottom) Day 2 at end-systole. SR-O and SR-R: Super-resolution using orthogonal and radial LA images, respectively. LR: low resolution

